# FOXC1 negatively regulates BMP-SMAD activity and Id1 expression during osteoblast differentiation

**DOI:** 10.1101/673889

**Authors:** Jordan. C. Caddy, Leiah. M. Luoma, Fred. B. Berry

## Abstract

Bone morphogenetic proteins regulate a diverse range of biological processes through their activation of SMAD1, 5, or 8 proteins that in turn regulate gene expression. These SMAD transcription factors achieve a layer of functional specificity in different cells types largely through actions with additional transcriptional regulatory molecules. In this report we demonstrate that the Forkhead Box C1 (FOXC1) transcription factor can modulate BMP signalling to impair expression of BMP4-responsive genes and prevent efficient osteoblast differentiation. We demonstrate that repression occurs downstream of BMP signalling and impacts the ability SMAD1 or 5 to activate gene expression. Repression of SMAD activity requires FOXC1 DNA-binding capacity and the transcriptional inhibitory domain of FOXC1. We report that FOXC1 inhibits BMP4 induction of *Id1* expression and identify a motif in the regulatory region of mouse Id1 gene that FOXC1 binds. We determine that this inhibition by FOXC1 binding does not affect SMAD1, 5, or 8 binding to its target sequence in the *Id1* gene. Finally we determine that elevated expression of FOXC1 can reduces expression osteogenic differentiation genes in mouse embryonic stems directed to the osteoblast lineage through BMP4 treatment. Together, these findings indicate that FOXC1 can negative regulate certain aspects of BMP4 signalling required for osteoblast differentiation. We propose that FOXC1 acts to attenuate the initial BMP-activated pathways that establishes osteoblast differentiation and allow for terminal osteoblast differentiation to conclude.

## 1. Introduction

Cell signalling events coupled with gene expression changes are vital for the coordinated control of cell differentiation and development. The bone morphogenetic protein (BMP) family of signalling factors participate in a number of important developmental pathways. BMPs were discovered based on their osteoinductive ability and thus have a crucial role in the formation of the skeleton [Lowery and Rosen, 2018; Nishimura et al., 2012; Urist, 1965]. However, BMP activity is required for the formation and differentiation of diverse cell types and organ systems outside of the skeleton, including the nervous system [Allan et al., 2003], vascular system [Goumans et al., 2018], and gastrointestinal system [Walton et al., 2016], among many others. Aberrant BMP signalling also contributes to many human disease phenotypes, including those affecting the musculoskeletal and cardiac systems, as well as many cancers [Salazar et al., 2016; Wang et al., 2014]. Thus it is evident that BMPs function in diverse range of cell types and biological processes. How the precise functional specificity is achieved by BMP signalling in different cell types is an active area of research.

BMPs are secreted molecules that dimerize and bind to a heterotetramer of BMP Receptor I and II (BMPRI and BMPRII) [Gomez-Puerto et al., 2019; Kretzschmar et al., 1997]. When bound by BMP ligand, BMPRII phosphorylates BMPRI, which in turn phosphorylates SMAD1, 5, or 8 proteins that facilitates binding to SMAD4 [Kretzschmar et al., 1997; Salazar et al., 2016; Zhang et al., 1997]. The SMAD complex then enters the nucleus and can regulate expression of target genes including *Id1, Runx2, Hey1, Dlx3, Col1a1*, and *Msx2* in bone forming cells [Hassan et al., 2004, 2006; Katagiri et al., 1994; Lee et al., 2000, 2003, 2006; Park and Morasso, 2002].

SMAD proteins are essential regulators of BMP signalling. However these factors bind to target DNA with low affinities to degenerate sequences in the genome [Chai et al., 2015; Johnson et al., 1999; Massagué et al., 2005; Shi et al., 1998]. Thus these proteins alone may not support cell and tissues specific BMP functions. Additional transcription factors work alongside SMAD proteins to enhance their functional fidelity in different cell types. For example, RUNX2 acts with SMAD1 or 5 to promote expression of *ColX* in chondrocytes. Members of the Forkhead Box (FOX) family of transcription factors have been shown to function along with SMAD proteins that are regulated by TGFβ signalling pathways. FOXH1 can physically interact with SMAD2 to modulate nodal activity [Labbé et al., 1998; Saijoh et al., 2000], and FOXG1 can regulate TGFβ signalling to control cell cycle progression in glioma cells [Seoane et al., 2004].

There is evidence to link the function of the FOXC transcription factors (which include FOXC1 and FOXC2 in mammals) to BMP signalling pathways. Primary micromass cultures from *Foxc1*^*−/−*^ fail to induce chondrogenic nodules in response to BMP2 treatment [Kume et al., 1998] and *Foxc1* function is required for activation of BMP-responsive genes in the mouse calvarium [Rice et al., 2003]. We have previously demonstrated that *Foxc1* is a target for BMP induced gene expression during early stages of osteoblast differentiation of mesenchymal progenitor cells [Hopkins et al., 2016]. *Foxc1* mRNA levels increased in BMP4-treated C2C12 cells through the direct binding of SMAD factors to the *Foxc1* regulatory region. Moreover, the enforced expression of *Foxc1* in C2C12 cells induced expression of early markers of osteoblast differentiation. Given these findings, we wished to further investigate the relationship between Foxc1 activity and BMP signaling in the regulation of osteoblast differentiation.

## 2. Methods

### 2.1. Tissue Culture

U2OS human osteosarcoma and C2C12 mouse myoblast cells were purchased from ATCC and were grown in Dulbecco’s Modified Eagle Medium (DMEM) (Sigma-Aldrich, St. Louis, MI Cat. # D6429) supplemented with 10% Fetal Bovine Serum (FBS). C2C12-Babe and C2C12-Foxc1 are cells stably transduced with empty pBABE or *Foxc1* as described previously [Mirzayans et al., 2012]. Mouse embryonic stem cells containing a tet-repressible *Foxc1* expression cassette [Correa-Cerro et al., 2011] were provided by T. Kume (Northwestern University). The cells were grown on gelatin coated tissue culture plates in DMEM containing 15% FBS, L-glutamine, MEM non-essential amino acids, Leukemia inhibitory factor (1000 U/ml), pen-strep, and doxycycline (1μg/ml) to suppress *Foxc1* expression.

### 2.2. Transactivating Assays

U2OS were seeded in 24 well plates at a density of 4× 10^4^cells per well. After 24 hours the cells were transfected with 150 ng/mL of each effector plasmid (FOXC1 wildtype (WT) or deletion/mutation constructs, empty vector (EV) or SMADs), 10-100 ng/mL of reporter plasmid (BRE-luc), 0.1 ng/mL of Renilla plasmid, and 3:1 volume:mass ratio of Mirus *Trans*IT®-LT1 transfection reagent (Mirus Bio LLC, Madison WI) to DNA. FOXC1-S131L and deletion constructs were described previously [Berry et al., 2002; Saleem et al., 2003]. Recombinant human BMP4 (50 ng/ml; R&D Systems, Minneapolis, MN) or equivalent volume of 4mM HCl (solvent for BMP4 reconstitution) was added 24 hours post-transfection and cells were incubated for an additional 24 hours. Luciferase assays were performed as recommended by the manufacturer (Promega, Madison WI). Each transfection consisted of three technical replicates and were performed at least three times.

### 2.3. Differentiation Assays

For osteoblast differentiation assays, 6-well or 35 mm plates were seeded with C2C12-pBABE, or C2C12-*FOXC1* at a density of 2×10^5^ cells/well. After 48 hours cells were washed with PBS, and treated with either BMP4 (50 ng/mL) or equivalent volume of 4 mM HCl in serum-reduced media (DMEM + 0.2% FBS). Cells were then fixed with 10 % formalin. Cells were then washed twice with PBS and stained using the BCIP/NBT liquid substrate system (Sigma-Aldrich) in the dark (covered with foil) for 3.5 hours. Staining solution was removed and the cells rinsed extensively in ddH_2_O before being imaged. Mouse ES cells (mES) were induced as embryoid bodies as described in [Kawaguchi et al., 2005].

### 2.4. RNA Extraction and Qualitative Reverse Transcriptase Polymerase Chain Reaction (qRT-PCR)

RNA extraction was carried out using the RNeasy Plus Mini Kit Animal Cell Spin technique (Cat# 74106 Qiagen, Mississauga, Canada) using the protocol provided by the manufacturer. RNA was then quantified using NanoDrop ND-1000 Spectrophotometer (Thermo Fisher Scientific, Waltham, MA). QuantiTect Reverse Transcriptase Kit (Qiagen Cat. # 205313) was used to create cDNA from 500 ng of RNA. 13 µL of 1:25 diluted cDNA was added to 16.25 µL KAPA 2x SYBR Fast Master Mix (KAPA Biosystems, Wilmington, MA) and 3.25 µL primer sets (Table 2.3) to create the master mix. The master mix was then vortexed briefly, spun down and 10 µL placed into each of three wells in a 96-well plate for technical replicates. Data were analysed using Bio-Rad CFX Manager version 3.0.125.0601 normalized to three housekeeping genes: *Gapdh, Hprt,* and *Actin B.*

### 2.5. Electrophoretic Mobility Shift Assay

Oligonucletode probes for the Id1 putative FOXC1 binding site labelled with the IRDye700 were obtained from Integrated DNA Technologies (Coralville, Iowa). Labelled sense and antisense probes were mixed at a final concentration of 4.5pmol/µL in 10 mM Tris, 20 mM NaCl and 50mM MgCl2. Nuclear lysates were pre-incubated for 10 minutes at room temperature in 10 mM HEPES, pH 7.9, 5% glycerol, 25 mM NaCl, 2 mM DTT, 0.1 mM EDTA and 0.05 µg/µL of poly dI-dC. Following pre-incubation, 1 pmol of labeled probe was added per reaction. For controls, 45 pmol of cold probe was added to reactions, or 1 µg of FOXC1 antibody as indicated. Binding reactions were performed at 25°C, in the dark. Reactions were resolved through a 5% polyacrylamide 1X TGE gels at room temperature using chilled 1X TGE in the dark, and visualized on a LI-COR Odyssey scanner (LI-COR, Lincoln, Nebraska).

### 2.6. Data analysis

All statistical analysis was performed using SigmaPlot 13. One-way ANOVA with Holm-Sidek post hoc or Dunn’s test were used.

## 3. Results

### 3.1. FOXC1 inhibits BMP-SMAD transcriptional regulatory activity

We first investigated whether FOXC1 could modulate BMP-SMAD activity using the well characterized BMP responsive BRE-Luc reporter vector [Korchynskyi and ten Dijke, 2002]. When U2OS cells were transfected with FOXC1 expression vectors along with BRE-luc and were treated with BMP4 (50 ng/ml) for 24 hours, we observed that activation of the BRE-Luc reporter was impaired by FOXC1. This inhibition was also observed in the presence of the paralogous FOXC2 protein (Figure 1A) and was also observed when cells were treated with BMP2 (data not shown). Transfection of FOXC1 and FOXC2 had no effect on phosphorylation of SMAD1,5,8 suggesting this inhibitory effect of FOXC1 and FOXC2 was occurring intracellularly following BMP induction (data not shown). We then examined whether FOXC1 was acting on the BMP-regulated SMAD proteins. Elevated expression of SMAD4 and SMAD5 alone could activate expression of BRE-luc in the absence of exogenous BMP4. When FOXC1 was co-expressed, this activation was diminished (Figure 1B). We were unable, however, to detect any potential protein-protein interactions occurring between FOXC1, SMAD4 or SMAD5 (data not shown). To further elucidate the mechanisms of how FOXC1 may impair BMP-SMAD activity, we used previously described FOXC1 deletion constructs [Berry et al., 2002] to identify what regions of FOXC1 were responsible this inhibition. As indicated in Figure 1C, we observed that removal of the FOXC1 N-terminal activation domain (AD1; FOXC1 29-551) partially restored some activation of BRE-Luc by BMP4, while removal of the FOXC1 C-terminal activation domain (AD2; FOXC1 1-366) had no effect on the FOXC1-mediated inhibition of the BRE-luc. When the C-terminus of FOXC1 was further deleted to remove the inhibitory domain (ID) of FOXC1(Δ216-553) or when only the Forkhead box domain (FHD) was expressed any inhibition of the BRE-luc by FOXC1 was lost. We instead observed elevated levels of BRE-luc activity. This elevation appeared to be unresponsive to BMP4 treatment. Finally we observed that FOXC1 inhibition of the BRE-luc activity was lost when we expressed FOXC1 containing a S131L mutation in the Forkhead domain that prevents FOXC1 DNA binding. Together these findings suggests elevated FOXC1 levels could prevent BMP4 activation of the BRE-luc reporter and this inhibition required FOXC1 inhibitory domain and DNA-binding activity.

**Figure 1.**
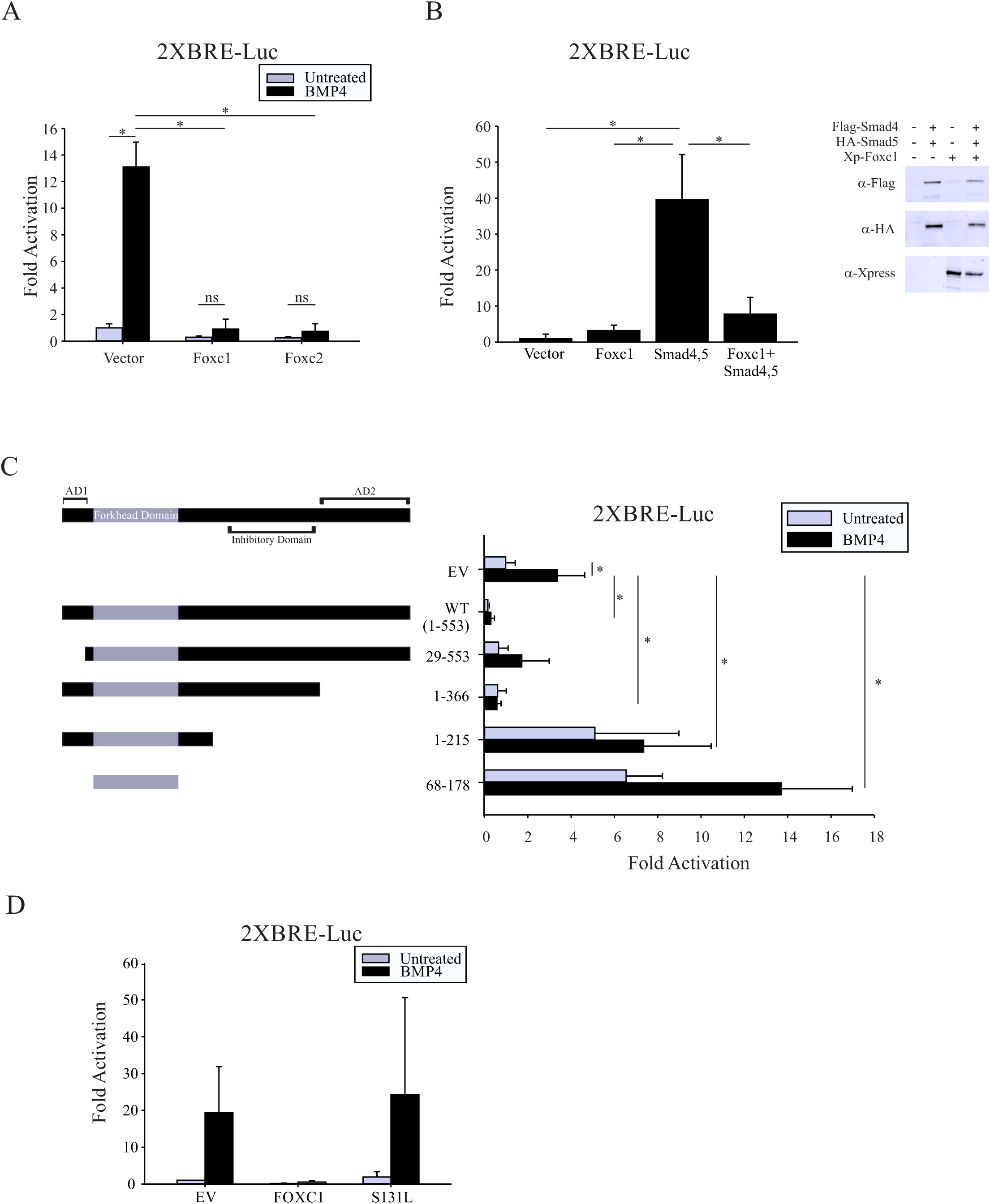
FOXC1 inhibits BMP4-induced transcriptional activation. A. U2OS cells were transfected with 2XBRE-luciferase reporter along with empty expression vectors or vectors expressing *FOXC1* or *FOXC2.* Cells were treated with BMP4 (50 ng/ml) 24 h post transfection. B. U2OS cells were transfected with vectors expressing combinations of Xpress(Xp) tagged FOXC1, Flag-SMAD4 and HA-SMAD5 along with 2XBRE luciferase. Immunoblot analysis confirmed expression differences in SMAD4 and SMAD5 protein levels did not account for inhibited activation of the 2XBRE in response to FOXC1. C. U2OS cells were transfected with full-length or truncated FOXC1 expression constructs along with 2XBRE-Luc. BMP4 (50 ng/ml) was added 24 h following transfection. All FOXC1 expression constructs contained nuclear localization signals found in the Forkhead Domain. (AD1= Activation Domain 1; AD2=Activation Domain 2). D. U2OS cells transfected with a wild-type FOXC1 or mutant FOXC1 (S131L) along with 2XBRE-luc. All transactivation assays were transfected in triplicate and each experiment was performed at least three times. Error bars represent the standard deviation. Asterisk indicate p<0.05. ns= non significant.

### 3.2. Elevated FOXC1 levels suppresses BMP4 induced osteoblast differentiation of C2C12 cells

We next examined whether elevated expression of FOXC1 impacted BMP signalling in a functional cellular context. C2C12 myoblast cells treated with BMP 2 or 4 trans-differentiate to osteoblast cells, as indicated by expression of osteoblast marker genes including Alkaline phosphatase (ALP) and Runx2 [Katagiri et al., 1994]. We previously demonstrated that stable, elevated expression of FOXC1 in C2C12 myoblast cells could ectopically induce ALP activity and expression of osteoblast genes in the absence of BMP signalling [Mirzayans et al., 2012]. C2C12-Foxc1 and control C2C12-Babe cells were cultured in growth media with or without BMP4 (50 ng/ml) and cells were stained for ALP activity as an indicator of osteoblast differentiation. In these experiments, a single treatment of BMP4 (50 ng/mL) was given at day 0 and cells were differentiated for a further 6 days. We confirmed elevated FOXC1 levels in the C2C12-FOXC1 cells (Figure 2B) and, in the absence of BMP4, C2C12-FOXC1 cells produced robust ALP activity after 6 days in culture (Figure 2A). However when these cells were treated with BMP4, little ALP activity was observed over the entire time course (Figure 2A). In contrast, the control cells produced little ALP activity in the absence of BMP treatment, while BMP addition led to an increase in ALP staining by 1 day that persisted over the 6 day period. We then monitored phosphorylation of SMAD1,5,8 by BMP4 and found this was not prevented in C2C12-FOXC1 cells (Figure 2C).

**Figure 2.**
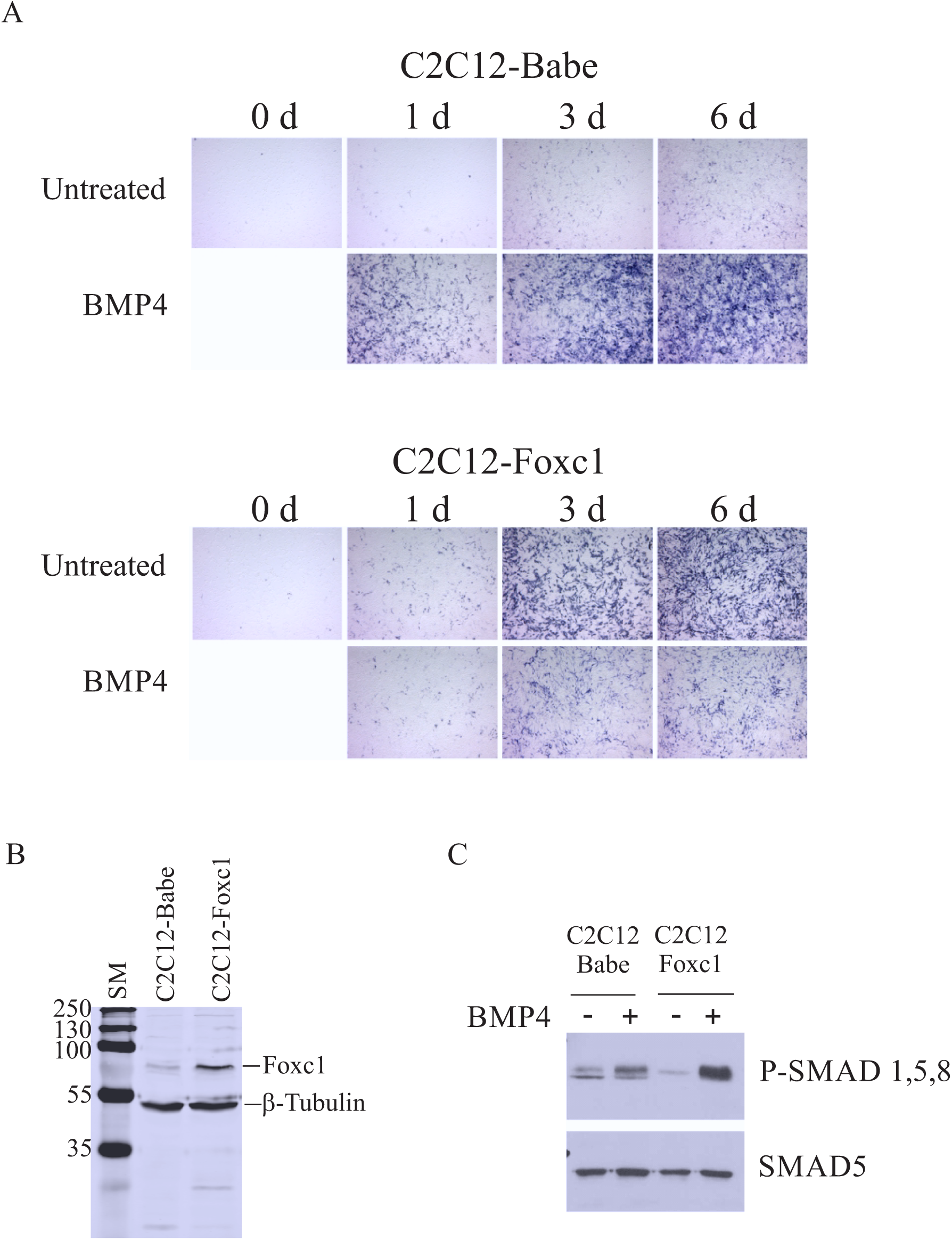
Elevated FOXC1 levels impairs BMP4-induced osteoblast transdifferentiation of C2C12 cells. A. C2C12 cells stably expressing FOXC1 (C2C12-FOXC1) or empty vector control cells (C2C12-Babe; [Mirzayans et al 2011]) were treated with or without BMP4 (50 ng/ml) for 6 days. Cells were fixed at the indicated time point and assayed for alkaline phosphatase activity. B. Immunoblot detection of FOXC1 indicates elevated protein levels in C2C12-Foxc1 cells compared to control C2C12-Babe cells. C. SMAD1,5 & 8 proteins were phosphorylated by BMP4 treatment in both C2C12-Babe and C2C12-Foxc1 cells.

Next, we assessed expression of BMP-induced osteoblast differentiation genes over this 6 day time course to determine whether elevated FOXC1 levels altered their gene expression patterns. Cells were treated with BMP4 as described above. *Id1, Runx2, Dlx3* and *Hey1* expression levels were increased in C2C12-Babe following 24 hours of BMP4 treatment (Figure 3). In contrast, levels of *Id1, Runx2* and *Hey1* mRNA in C2C12-FOXC1 cells were reduced at 24 hours of BMP4 treatment compared to control C2C12-BABE cells (Figure 3). These data suggest that elevated FOXC1 can impair BMP4 induced osteoblast trans-differentiation of C2C12 cells.

**Figure 3.**
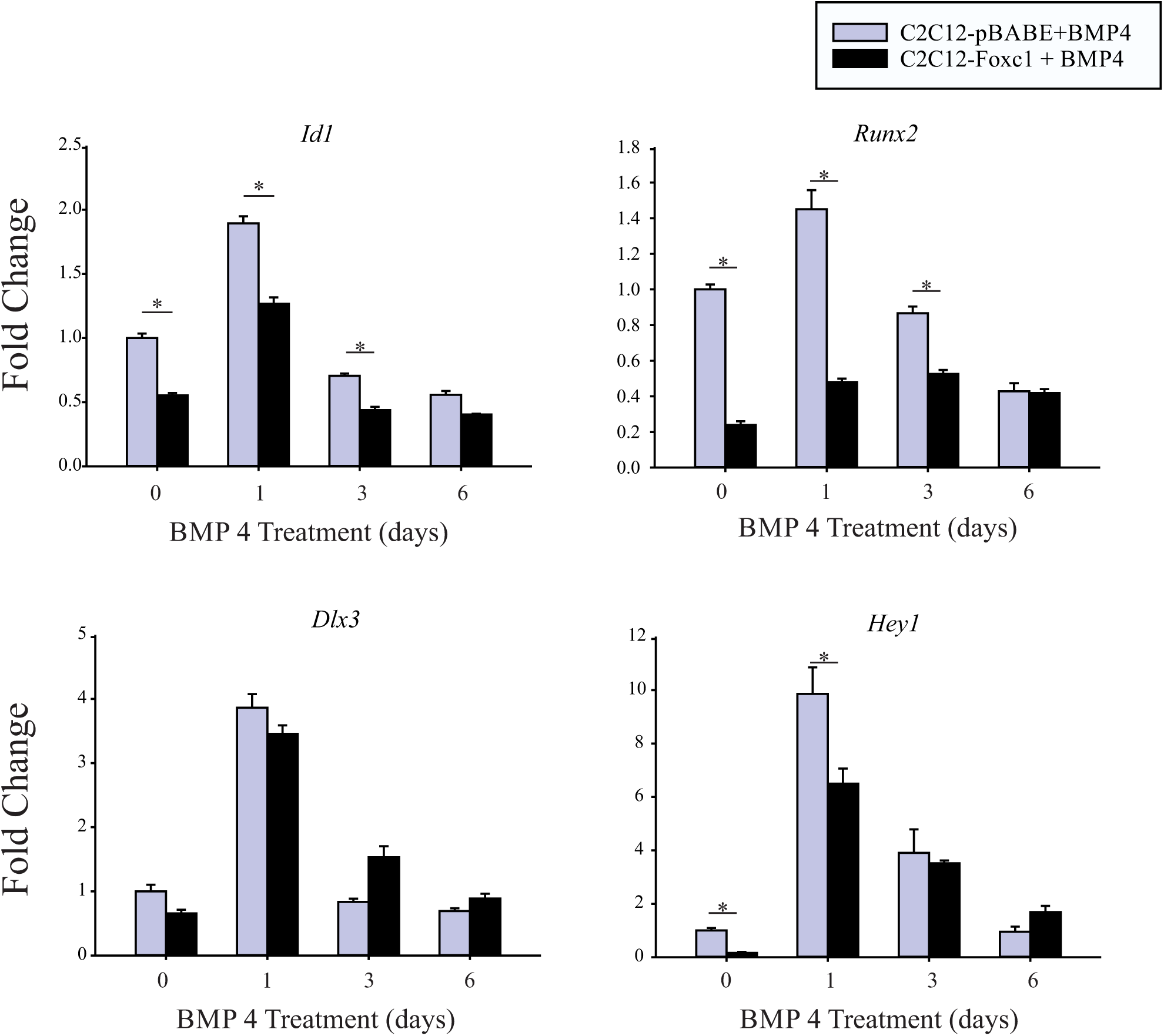
Expression of BMP-induced osteoblast differentiation genes is reduced in response to elevated FOXC1 levels. C2C12-Babe and C2C12-Foxc1 cells were treated with BMP4 (50 ng/ml) and RNA was harvested at the indicated time points. Error bars represent standard error of the mean. Asterisk indicate p<0.05. Experiments were performed on three biological replicates.

### 3.3. Foxc1 directly inhibits *Id1* expression

Since the responsive element in the BRE-luc reporter used in Figure 1 was derived from the mouse *Id1* gene, we explored further the potential direct regulation of *Id1* expression by Foxc1. We observed an induction of *Id1* mRNA expression in the parental C2C12, C2C12-BABE and C2C12-FOXC1 cells following 24 h BMP4 treatment (Figure 4A). However, expression levels of *Id1* mRNA were lower in C2C12-FOXC1 in both untreated and 24 h BMP treated cells. We then employed mouse embryonic stem cell line [Nishiyama et al., 2009] carrying a Tet-inducible Flag-Foxc1transgene to determine whether an acute activation of *Foxc1* expression could affect induction of *Id1* mRNA by BMP4 treatment. When Foxc1 levels were activated, we did observe that *Id1* mRNA induction by BMP4 treatment was blocked (Figure 4B). We identified a putative Foxc1 binding motif (GTAAATAAA) in the regulatory region of *Id1* gene, located between the BRE and the TATA box (Figure 4C). We used *Id1* promoter deletions [Tournay and Benezra, 1996] to test whether FOXC could regulate *Id1* activity. We found that FOXC1 could prevent the induction of *Id1* expression by BMP4 in the 1.5 kb reporter containing the BRE and Foxc1 binding sites. Deletion of the BRE prevented any activation of reporter genes by BMP4 whether the FOXC1 element was present or not. We then determined that Foxc1 could in fact bind to the putative FOXC1 binding element in *Id1* by Electrophoretic Mobility Shift Assays (EMSA) (Figure 4D). FOXC1-DNA interactions could be competed off by treating with Foxc1 antibodies and with an excess of unlabelled probe from the *Id1* promoter (Id1) or from a well characterized Foxc1 binding motif (BS) [Saleem et al., 2001]. The addition of SMAD4 along with SMAD1 or 5, had no effect on FOXC1 binding to the *Id1* promoter, nor did it promote the formation of a higher order complex with FOXC1. Finally, we assessed whether elevated FOXC1 levels prevented SMAD1, 5 or 8 binding to the BRE in the *Id1* promoter. We performed chromatin immunoprecipitation assays (ChIP) and verified that BMP-responsive SMAD proteins were still able to bind to the *Id1* BRE following BMP4 treatment in C2C12-FOXC1 cells (Figure 5).

**Figure 4.**
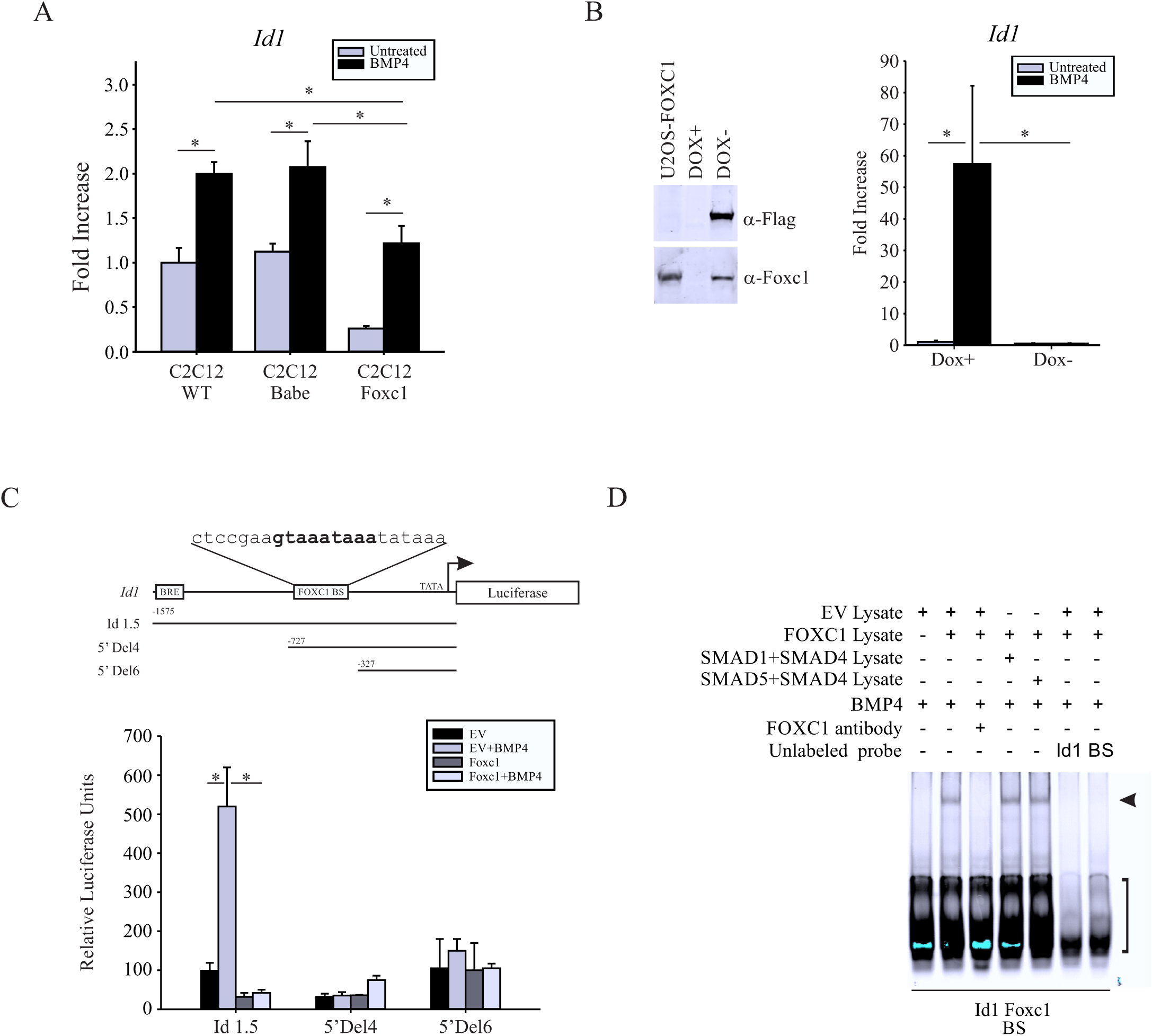
*ID1* is a directed target of FOXC1 expression. A. Expression of *Id1* mRNA was detected in parental C2C12, C2C12-Babe or C2C12-Foxc1 cells following 24 hours BMP4 (50 ng/ml) treatment by qRT-PCR. B. Mouse embryonic stem cells containing a Tet-off, Foxc1-inducible gene were treated with or without doxycycline (Dox) for 48 hours. Induction of Foxc1 was confirmed by immunoblotting with Foxc1 and Flag antibodies. Expression of *Id1* mRNA was assessed by qRT-PCR. C. A FOXC1 consensus binding site (bold) was found in the mouse Id1 promoter between the BRE and transcription start site. U2OS cells were transfected with 1.5 kb Id1 reporter, or with truncated reporters that retained (5’ Del4) or deleted (5’ Del6) the FOXC1 binding site (BS) along with empty vector (EV) or *Foxc1* expression vectors. Cells were treated with BMP4 24h post transfected and harvested 24 hours later. D. EMSA assays demonstrating FOXC1 binding to the Id1 promoter. Cell extracts from U2OS cells transfected with empty vector (EV), Foxc1, Smad1, Smad5 or Smad4 were incubated with labelled probe representing FOXC1 binding site of the Id1 promoter. As a control reactions were incubated with an excess of unlabelled probe from the Id1 gene or a well-established Foxc1 binding sequence (BS) [Saleem et al 2001]. Error bars represent the standard error of the mean (A,B) or the standard deviation (C). Asterisk indicate p<0.05. Arrowhead indicates Foxc1-DNA interaction. Square bracket indicates unbound probe.

**Figure 5.**
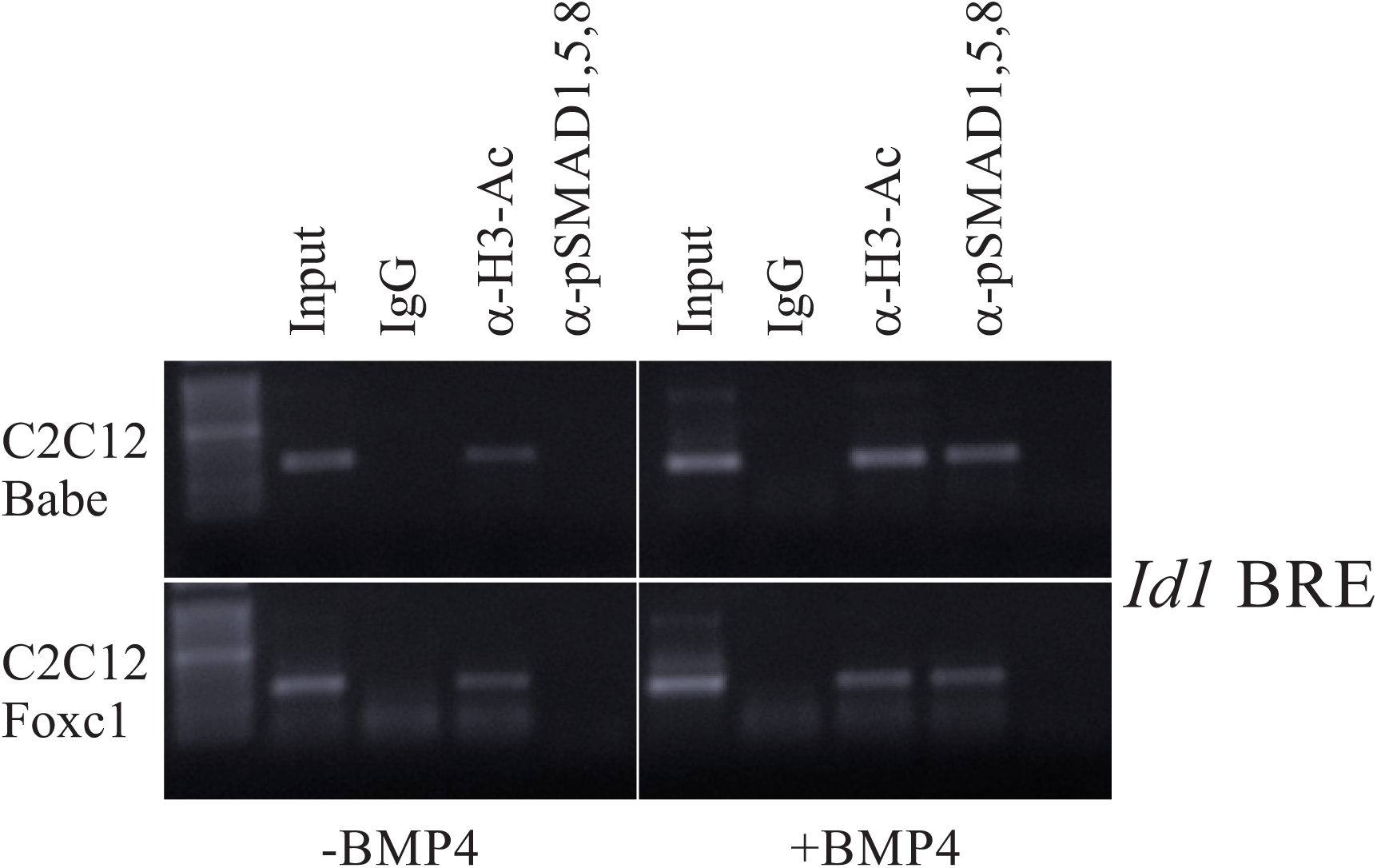
SMAD1,5 or 8 binding to the *Id1* BRE is not affected by elevated FOXC1 levels. Chromatin from C2C12-Babe or C2C12-Foxc1 treated with or without BMP4 (50 ng/ml; 24 h) was immunoprecipitated with antibodies recognizing acetylated Histone H3, phospho(p)-SMAD1,5,8 or with rabbit IgG as a negative control. Recovered DNA was then amplified by PCR using primers flanking the BRE of the mouse *Id1* gene.

### 3.4. Elevated Foxc1 levels prevented osteoblast differentiation of mouse ES cells

Although BMP4-induced differentiation of C2C12 is a well-established model of osteoblast differentiation, we wanted to examine the role of Foxc1 in modulating BMP4-regulated differentiation in an additional biological system. Mouse embryonic stem cells can be directed towards the osteoblast lineage by growing cells as embryoid bodies in the presence of Retinoic acid, followed by attached growth in the presence of BMP4 ([Kawaguchi et al., 2005] (Figure 6A). We used mouse embryonic stem cells engineered to express Foxc1 in a tetracycline repressible system [Nishiyama et al., 2009]. Removal of the tetracycline analog, doxycycline, resulted in a robust activation of *Foxc1* expression levels (Figure 6B). When mES cells were induced to differentiate toward the osteoblast lineage through BMP4 treatment in the presence of Dox for 15 days, we observed elevated expression of osteogenic genes *Runx2, Col1a1 Sp7 (Osterix)* and *Bglap (Osteocalcin)* compared to cells at Day 0. When *Foxc1* expression was induced, we observed a decrease in expression of *Col1a1, Sp7* and *Bglap* mRNA compared to uninduced cells. These findings suggest that prolonged expression of elevated expression of FOXC1 in mES cells can reduce the efficiency of BMP4-mediated osteoblast differentiation.

**Figure 6.**
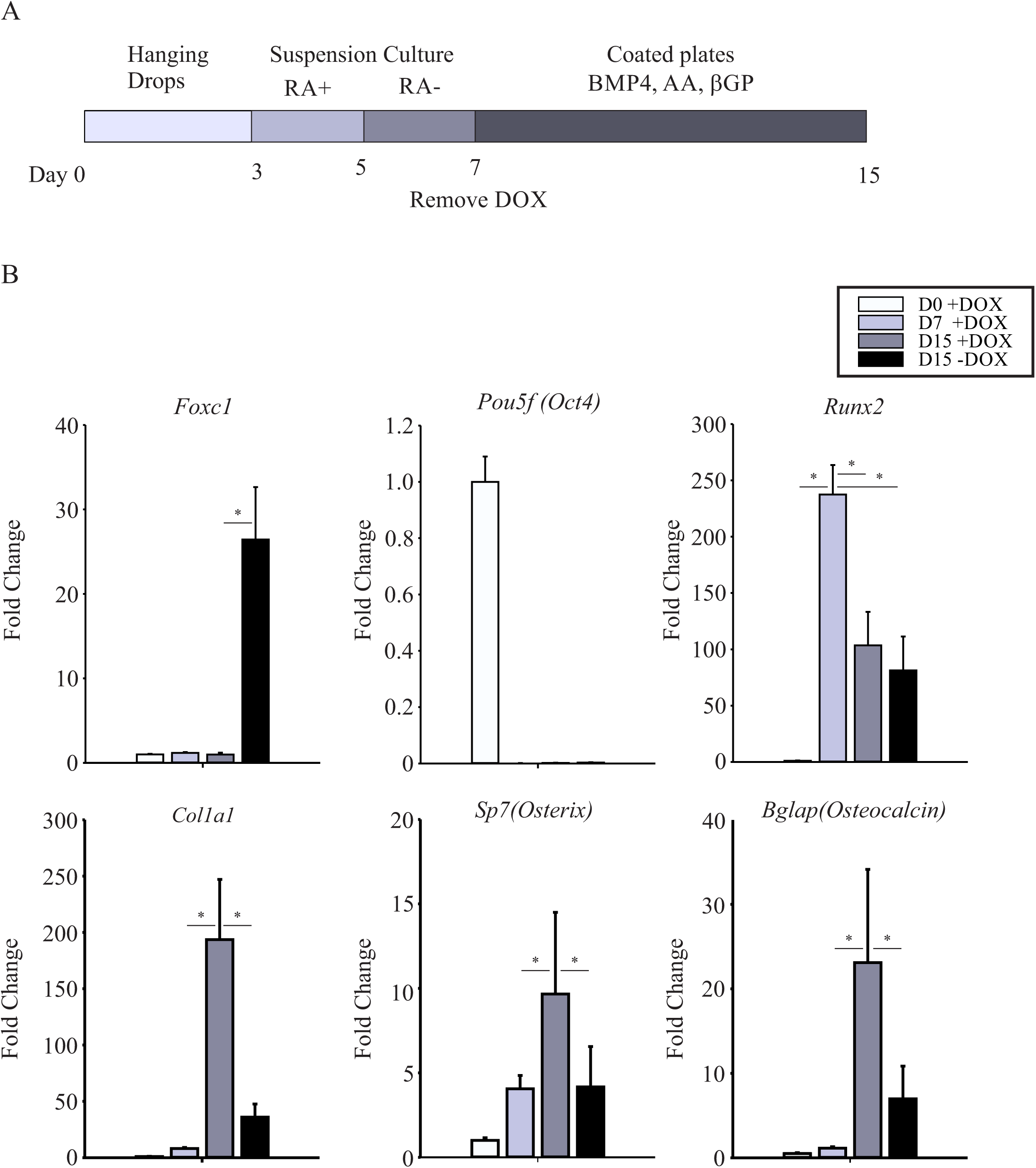
Elevelated FOXC1 levels impairs BMP4-induced osteoblast differentiation of mouse embryonic stem cells. A. Outline of the osteoblast differentiation procedure of mouse ES cells [Kawaguchi J et al., 2005]. Cells were grown as hanging in LIF-deficient media for 3 days, then transferred to suspension culture and grown for 2 days with Retinoic acid (10^−7^ M) followed by 2 days without. On day 7, cells were cultured under adherent conditions and were treated with osteoinductive media (BMP4 (100 ng/ml), Ascorbic acid (AA; 50 μg/ml) and β-glycerophosphate (βGP; 50 mM)) for an additional 8 days. Doxycycline was removed on Day 7. B. Expression levels of osteogenic differentiation genes analysed by qRT-PCR. Error bars represent standard error of the mean. Asterisks indicate p<0.05. qRT-PCR-reactions were performed in triplicate on three biological replicates.

## 4. Discussion

In this report, we demonstrated that elevated *Foxc1* levels can prevent BMP-SMAD transcriptional regulatory activity and inhibit BMP4-induced activation of *Id1* gene expression. The BRE from the *Id1* gene is a well-established and characterized motif to monitor BMP-regulated SMAD activity [Katagiri et al., 2002; Korchynskyi and ten Dijke, 2002]. We found that Foxc1 prevented activation of this reporter by BMP4 treatment and when cells over-expressed SMAD1 and 4. Moreover, these data and our observation that SMAD1,5,8 proteins were phosphorylated by BMP4 in the presence of elevated FOXC1 levels, suggests that FOXC1 does not act to prevent the propagation of BMP4 signalling to SMAD proteins but rather may act to inhibit SMAD transactivation of BMP4-responsive targets. We were unable to detect any potential protein-protein interactions occurring between FOXC1, SMAD4 or SMAD5 (data not shown). We also demonstrated that Foxc1 does not prevent binding of SMAD1 protein to the BRE element, suggesting FOXC1 inhibition does not occur in a competitive manner. The use of Foxc1 deletion constructs revealed that DNA-binding activity and the Foxc1 inhibitory domain were needed to prevent activation of the BRE by BMP4. The inhibitory domain of FOXC1 has been shown to be required for negative regulation of genes by FOXC1. How this region achieves this activity has yet to be determined. Putative binding motif for Groucho/TLE transcriptional repressors are present in the inhibitory domain of FOXC1, as well as in other FOX proteins [Yaklichkin et al., 2007]. We were unable to detect any binding of TLE proteins with FOXC1 and co-transfection of TLE1, TLE3 or TLE4 along with FOXC1 did not augment any transcriptional inhibition of the BRE by FOXC1 (data not shown).

We observed that elevated Foxc1 expression in prevented activation of *Id1* mRNA expression in response to BMP4 treatment in C2C12 cells, suggesting *Id1* mRNA expression is regulated by FOXC1. Moreover, we determined that FOXC1 could prevent activation of an *Id1* promoter reporter containing the BRE motif [Tournay and Benezra, 1996]. *We also identified a Forkhead box binding motif in the Id1* regulatory region that was bound by FOXC1 protein and was required for inhibition of BMP4 signalling by FOXC1. Together these findings suggest that FOXC1 acts as a negative regulator of *Id1* expression by BMP4 signalling. *Id1* gene expression is also activated by BMP signalling during the initiation osteoblast differentiation and *Id1* expression is reduced as differentiation proceeds [Peng et al., 2004]. Like Foxc1, enforced expression of *Id1* prevented osteoblast differentiation [Peng et al., 2004]. Thus, we propose that Foxc1 acts as a negative regulator of the BMP signalling pathway to extinguish expression of genes following their initial activation by BMP-signalling at early stages of differentiation. *Msx2* activation by BMP was prevented by Foxc1 activity, further supporting a role for Foxc1 to negatively influencing expression genes activated by BMP [Sun et al., 2013]. Clearly the identification of additional BMP-regulated genes controlled by FOXC1 will need to be determined.

Our previous research showed that Foxc1 expression was differentially regulated by BMP signalling at different stages of osteoblast differentiation [Hopkins et al., 2016]. Foxc1 expression is induced by BMP4 in C2C12 cells and is required for BMP4-induced osteoblast differentiation of these cells. However, expression of Foxc1 was reduced by BMP4 treatment in committed MC3T3 osteoblasts as they terminally differentiated, suggesting Foxc1 may act in the specification of the osteoblast lineage but is not required for terminal differentiation. Our observation in this study that elevated expression of *Foxc1* in C2C12 cells prevented BMP-induced osteoblast differentiation supports this notion. Recently, elevated expression of *Foxc1* in MC3T3 cells was shown to accelerate osteoblast differentiation independent of BMP-signalling [Shen et al., 2019]. FOXC1 inhibitory effects on BMP-induced osteoblast differentiation of dental pulp have also been observed [Xiao et al., 2018]. These data, along with our finding that enforced expression of Foxc1 in mouse embryonic stem cells reduced the osteoblast differentiation following BMP4 treatment further supports the idea that *Foxc1* has inhibitory effects on osteogenic differentiation. Loss of Foxc1 function mice prevents the formation of the calvarial bones [Kume et al., 1998]. In the skulls of Foxc1-deficient mice, BMP4 signaling is expanded where rudimentary bones form and ectopic ossification, along with expanded expression of osteogenic marker genes, is observed [Sun et al., 2013]. In the jaws of Foxc1 deficient mice, sygnathia, a bony fusion of the upper and lower jaws is observed, along with ectopic ossification of osteoblast progenitors in the jaw primordium [Inman et al., 2013], further suggesting a role for FOXC1 in regulating the timing of osteoblast differentiation. We propose that BMP-induced osteoblast differentiation requires Foxc1 function for the initial commitment of mesenchymal progenitors to the osteoblast lineage by BMPs but its expression of Foxc1 is downregulated as terminal differentiation is completed.

*Id1* was identified as an inhibitor of myogenic differentiation. ID1 protein fulfils this role by binding to and sequestering E2A proteins from basic helix-loop-helix transcription factors, such as MYOD [Jen et al., 1992]. Given that BMP2 treatment of C2C12 cells prevents myogenic differentiation and promotes osteogenic differentiation, it was proposed that upregulation of *Id1* expression account for this differentiation fate switch [Katagiri et al., 2002]. The role of ID1 in this process may extend beyond simply suppressing the myogenic phenotype and may have a role in regulating osteogenic differentiation. However, similar to FOXC1, ID1 function may regulate specific stages of osteogenesis. Compound heterozygous Id1^+/−^;Id3^+/−^ mice display reduced cranial suture formation [Maeda et al., 2004].While Id1−/− mice are viable and have a reduced bone mass, however this is likely the result of accelerated osteoclast function as expression of osteoclast-forming genes are elevated Id1−/− mice [Chan et al., 2009]. BMP-independent activation of Id1 expression by YAP1 is required for the promotion of MC3T3 osteoblast differentiation in response elevated Yap1 expression [Yang et al., 2019]. However, there are instances whereby Id1 and other Id gene function to inhibit osteogenic differentiation. CBAF1 DNA-binding and expression of osteogenic genes Alp and Ostecalcin is impared by direct binding with ID proteins. Moreover, a targeted down regulation of Id1 expression rescues osteogenic differentiation that was repressed from prolonged expression of *Bmp2* and *Vegf* [Song et al., 2011]. Although these findings all point to both positive and negative regulatory function of Id genes in osteogenic differentiation, the exact mechanism of how and when Id1 function to control this process will be an area of future interest.

Although much research has informed us on the cellular events that initiate a signalling response to factors such as BMPs, we know little regarding the mechanisms that terminate or damper these signals. During development, signalling cascades are not simply events that are activated and left on for the remainder of morphogenesis. Exquisite regulation by activator and inhibitors to fine-tune signaling output. For example BMP4 acts to prevent the sclerotome specification in the somites, but BMP4 is required at later stages of development for the same sclerotome cells to differentiate into chondrocytes [Murtaugh et al., 1999]. Moreover the differentiation of mES cells into mesoderm cell lineages is achieved through the carefully timed application of BMP signalling antagonists and agonists, further indicating the precise need to control this signalling pathway [Chal et al., 2015; Craft et al., 2015]. Our work suggests that FOXC1 is an additional factor that acts to fine tune the effect of BMP-signalling during osteoblast differentiation by downregulating expression of Id1.

## Acknowledgements

This work was supported by research funds awarded to F.B.B. from the Canadian Institutes of Health Research (MOP114921) and from the Edmonton Civic Employees Charitable trust fund. J.C.C was the recipient of Canada Graduate Scholarship awarded by the Natural Sciences and Engineering Research Council of Canada. F.B,B. holds the Shriners Hospital for Children Endowed Chair in Pediatric Scoliosis Research.

